# PRMT1 regulates EGFR and Wnt signaling pathways and is a promising target for combinatorial treatment of breast cancer

**DOI:** 10.1101/2021.10.22.465320

**Authors:** Samyuktha Suresh, Solène Huard, Amélie Brisson, Fariba Némati, Coralie Poulard, Mengliang Ye, Elise Martel, Cécile Reyes, David C. Silvestre, Didier Meseure, André Nicolas, David Gentien, Muriel Le Romancer, Didier Decaudin, Sergio Roman-Roman, Thierry Dubois

**Author notes:** <u>**Corresponding author:**</u> Thierry Dubois, Institut Curie - Centre de Recherche, Département de Recherche Translationnelle, Groupe “Biologie du Cancer du Sein”, 26 rue d’Ulm, 75005 Paris, France.

## Abstract

Identifying new therapeutic strategies for triple-negative breast cancer (TNBC) patients is a priority as these patients are highly prone to relapse after chemotherapy. Here, we found that protein arginine methyltransferase 1 (PRMT1) is highly expressed in all breast cancer subtypes. Its depletion decreases cell survival by inducing DNA damage and apoptosis in various breast cancer cell lines. Transcriptomic analysis and chromatin immunoprecipitation revealed that PRMT1 regulates the epidermal growth factor receptor (EGFR) and the Wnt signaling pathways, reported to be activated in TNBC. The enzymatic activity of PRMT1 is also required to stimulate the canonical Wnt pathway. Recently developed type I PRMT inhibitors decrease breast cancer cell proliferation and show anti-tumor activity in a TNBC xenograft model. These inhibitors display synergistic interactions with some chemotherapies used to treat TNBC patients, as well as the EGFR inhibitor, erlotinib. Therefore, targeting PRMT1 in combination with drugs used in the clinic may improve current treatments for TNBC patients.

**Significance:** This study highlights the requirement of PRMT1 for breast cancer cell survival and demonstrates the potential of targeting type I PRMTs in combination with chemotherapies in triple-negative breast cancer.

## Introduction

Breast cancer (BC) is a heterogeneous disease with molecularly distinct subtypes displaying different clinical outcomes and responses to therapies (1). Patients with “triple-negative” breast cancer (TNBC, lacking the expression of estrogen and progesterone receptors and Her2 overexpression) are mainly treated with conventional chemotherapies (1, 2). However, these patients have the worst prognosis as their treatment is challenging, owing to the inter- and intra-tumor heterogeneity leading to resistance to chemotherapy and relapse (2). Therefore, more efficacious treatments are needed to improve TNBC patient survival.

EGFR is overexpressed in more than 70% of TNBC patients and is associated with a metastatic phenotype (3). However, targeting this receptor as a monotherapy has shown only modest to low efficacy in clinical trials for TNBC patients (3). The Wnt signaling pathway is another pathway activated in TNBC through an overexpression of the transmembrane receptors, Frizzleds and co-receptors low-density lipoprotein receptor-related proteins (LRP6 and LRP5) (4–6). Wnt ligands (such as Wnt3a), which are secreted upon palmitoylation by the enzyme porcupine (7), activate the Wnt pathway by binding to the transmembrane receptors Frizzleds and co-receptors LRP5/LRP6. This initiates the release of β-catenin from the destruction complex including Dishevelled and Axin. Free β-catenin translocates into the nucleus and binds to the TCF/LEF family of transcription factors to activate the expression of Wnt target genes (4,5,7).

Arginine methylation of histones and non-histone proteins is a post-translational modification catalyzed by Protein Arginine Methyltransferases (PRMTs) (8–13). Substrate arginine can either be monomethylated or dimethylated (symmetrically or asymmetrically) by PRMTs. Asymmetric dimethylation is carried out by Type I PRMTs (PRMT1-4, PRMT6, and PRMT8) with PRMT1 being responsible for most of this modification (8–13). PRMTs are ubiquitously expressed, apart from PRMT8 which is brain-specific (12). Several PRMTs are overexpressed in various cancer types, including breast cancer (13–16) and are emerging as attractive therapeutic targets (9,10,12,14). Specific PRMT inhibitors have been recently developed and those against PRMT5 (10), the main type II PRMT, are being evaluated in Phase I clinical trials (NCT03573310, NCT03854227, NCT02783300). PRMT1-specific inhibitors are not yet available, nevertheless, two type I PRMT inhibitors (MS023, GSK3368715) have been developed showing more efficacy towards PRMT1, PRMT6 and PRMT8 (17, 18). GSK3368715 is in a Phase I clinical trial for diffuse large B-cell lymphomas and solid cancers (NCT03666988).

Arginine methylation regulates several cellular processes including transcriptional regulation and signal transduction (8,11,12). PRMT1 and PRMT5 regulate the EGFR signaling pathway by methylating EGFR (in colorectal and TNBC cells) (19–21), or by methylating histones on the EGFR promoter (in glioblastoma or colorectal cells) (22, 23) to regulate its transcription. Furthermore, some PRMTs regulate the canonical Wnt signaling pathway (12). Indeed, PRMT1 could either activate this pathway by methylating G3BP1 or G3BP2 (24, 25), or inhibit it by methylating Axin and Dishevelled (26, 27). Whether PRMT1 regulates the Wnt pathway in breast cancer cells is not known.

Among the different BC subtypes, PRMT1 has been mainly studied in luminal BC due to its well-described function as a transcriptional coactivator of estrogen receptor (ER) (14, 28). In this study, we found that PRMT1 is highly expressed in BC and that it is required for BC cell survival. We show that PRMT1 activates the EGFR and Wnt signaling pathways at the transcriptomic level. Type I PRMT inhibitors synergize with cisplatin, camptothecin, cyclophosphamide, and erlotinib, *in vitro*, and reduces tumor growth in a TNBC xenograft model. Together, our results suggest that combining type I PRMT inhibitors with drugs used in the clinic, could represent a promising therapeutic strategy for TNBC patients.

## Materials and methods

### Human samples and immunohistochemistry

Our cohort is composed of 35 luminal A (LA), 40 luminal B (LB), 46 TNBC, 33 Her2+, 18 normal breast tissues, and 14 TNBC cell lines (16, 29). DNA (Affymetrix SNP 6.0) and RNA (Affymetrix U133 plus 2.0) microarrays on this cohort have been described (29). Immunohistochemistry (IHC) using a PRMT1 antibody (Table S1) was performed as described (29).

### Cell culture, RNA interference, antibodies, small-molecule inhibitors, and primers

Cell lines were purchased from the American Type Culture Collection (ATCC), authenticated in 2021 by short tandem repeat profiling (data not shown), and cultured as described (16). The murine cell lines, L-cells and L-Wnt3a were obtained from Institut de Recherches Servier, France. The MDA-MB-231 cell line was a kind gift from Dr. Mina Bissell (University of California, Berkeley). References for antibodies, siRNAs, primers and drugs/small-molecule inhibitors are listed in Table S1.

### Cellular Assays

siRNA (20nM) transfection was performed using Interferin (Polyplus, 409-50), according to the manufacturer’s instructions. Cell proliferation was determined by MTT (Sigma, M2128-1G), WST-1 (Sigma-Aldrich, 11644807001) or CellTiterGlo (Promega, G7572) assays. Apoptotic activity was determined by the Caspase-Glo 3/7 luminescent assay (Promega, G8092), Annexin-V staining (Roche, 11988549001) or Western blot analysis. 2D colony formation and soft-agar assays were performed as described (16, 29). Chromatin immunoprecipitation (ChIP) was performed using the simple ChIP plus enzymatic chromatin IP Kit (Cell signaling, 9004), following manufacturer’s protocol. Wnt target gene expression analysis by RT-qPCR and β-catenin-activated reporter (BAR) luciferase assay were performed as described (30). Drug combinations were performed by treating cells with varying concentrations of the drugs/inhibitors (Table S1) and cell viability was measured by CellTiterGlo assay (Promega, G7572).

### PRMT1 Transcriptomic analysis

Transcriptomic analysis of PRMT1-depleted cells was performed using Affymetrix HTA 2.0 microarray. Significant differential gene expression between control and PRMT1 siRNA with an adjusted p-value cut-off of 0.05 was considered (Table S2). Gene enrichment pathway analysis was performed using the REACTOME database from GSEA website (31).

### GSK3368715 treatment in mice

Swiss-nude mice were purchased from Charles River and their care and housing were conformed to the French Ethical Committee guidelines. Tumors derived from MDA-MB-468 cells were grafted into control (n=6, vehicle-treated) and study (n=6, GSK3368715-treated) mice.

### Statistical analysis

R software and GraphPad Prism 7 were used for statistical analyses. Pearson or Spearman correlation were used to estimate an association between two variables. For cellular assays, p values were calculated using the Student’s t test, unless specified otherwise. Independency between tumor subtypes in the tumor microarray analysis was assessed using Fisher’s exact test.

### Data availability

The transcriptomic data generated in this study are available in supplementary data files (Table S2).

Additional and detailed experimental procedures are available in supplemental methods.

## Results

### PRMT1 is overexpressed in all the breast cancer subtypes compared to normal breast tissue

With the goal of identifying enzymes overexpressed in BC compared to normal tissue, we have performed gene expression profiling on a cohort of 154 human BC biopsies and healthy breast tissues (6,16,29). We found that PRMT1 mRNA is overexpressed in all BC subtypes compared to normal tissues, and observed the highest expression in TNBC (Figure 1A, left panel). The higher expression of PRMT1 mRNA in TNBC was confirmed in the publicly available database - the cancer genome atlas (TCGA) cohort (Figure 1A, right panel). We examined whether variations in PRMT1 expression could be a result of genomic alterations by analyzing DNA microarrays. Indeed, there was a correlation between PRMT1 mRNA and gene copy number within the whole cohort (Sup. Fig 1A). Interestingly, the PRMT1 locus showed significantly more gains in TNBC than the luminal BC subtypes and normal tissue (Figure 1B, Table S3). The PRMT1 mRNA levels also correlated positively with proliferation (Ki67 mRNA) in our cohort (Sup. Fig 1B).

**Figure 1.**
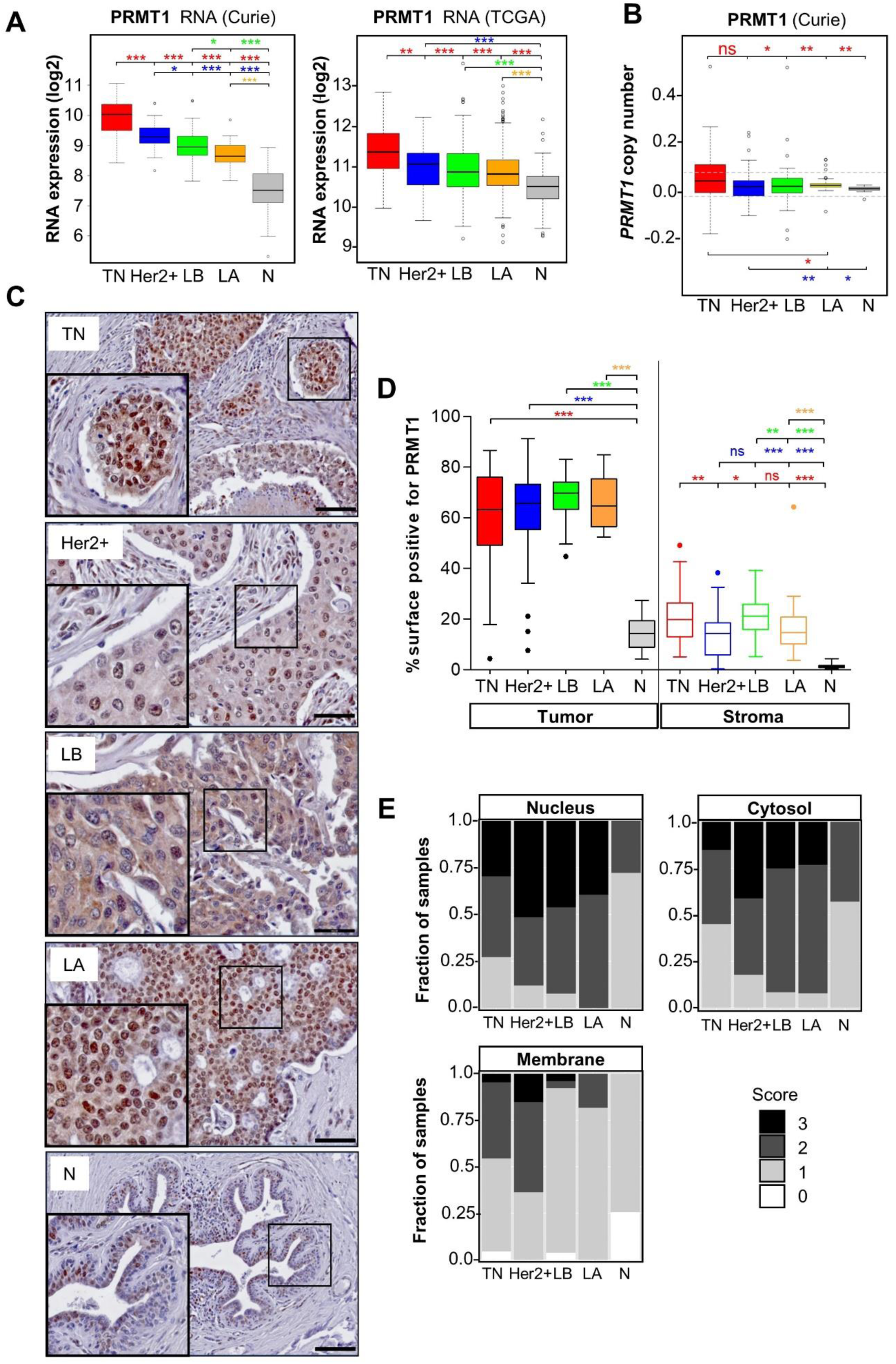
PRMT1 is highly expressed in breast tumors. **A,** Higher levels of PRMT1 mRNA in breast cancer. PRMT1 RNA expression in TNBC (TN, red), Her2+ (blue), Luminal B (LB, green), Luminal A (LA, orange), and healthy breast tissue (N, grey) in Curie (left panel) and TCGA (right panel) cohorts is illustrated by box plots (log2 transformed). **B,** Higher PRMT1 DNA copy number (CN) in TNBC in the Curie cohort. PRMT1 DNA CN determined by Affymetrix microarray analysis is presented in boxplots (smoothed segmented CN signal), with dashed lines indicating the thresholds retained to call CN gains and losses (see Table S3 for the number of samples showing loss or gains). **C,** Higher levels of PRMT1 protein in BC. PRMT1 protein levels were analyzed by IHC in the Curie cohort. A representative image of PRMT1 staining is shown for the different BC subtypes (scale bar = 50 μM). **D,** Quantification of the tumoral (left) or stromal (right) surface positive for PRMT1 staining represented as a percentage compared to the total surface. **E,** Quantification of PRMT1 staining in the different cellular compartments (0: no staining, 3: the strongest staining). *p < 0.05; **p<0.01; ***p < 0.001, as calculated using the Student t-test (A), Fischer exact test (B) or Mann Whitney test (D).

To understand the clinical significance of PRMT1 mRNA expression, we plotted survival outcomes from the KM-plotter database (www.kmplot.com) (32). High PRMT1 mRNA expression was associated with poor recurrence-free survival (RFS) in all BC (p=1×10^-8^, Sup. Fig 1C), as previously reported (33). However, this analysis does not take into account that PRMT1 is differentially expressed among the BC subtypes (Figure 1A) which are associated with different prognosis. Therefore, we performed this analysis within the different BC subtypes. High PRMT1 mRNA levels were associated with poor RFS in LA (p=2.5×10^-6^) and LB (p=0.007) (Sup. Fig 1C, top panel). Although this trend was seen in the Her2+ subtype, it was not statistically significant (p=0.13) (Sup. Fig 1C, bottom left panel). Conversely, high PRMT1 mRNA expression showed better RFS (p=0.02) within the TNBC subtype (Sup. Fig 1C, bottom right panel).

As mRNA and protein levels do not always coincide, we studied PRMT1 protein expression in BC and normal tumors using a commercial PRMT1 antibody. We first validated this antibody for immunohistochemistry (IHC) staining in a TNBC cell line (MDA-MB-468) fixed in the same method as the tissue samples (Sup. Fig 1D). IHC analysis confirmed that PRMT1 is highly expressed in all BC subtypes compared to normal tissue (Figure 1C, 1D). In contrast to mRNA expression, we did not observe any significant difference in PRMT1 protein expression level between the different BC subtypes (Figure 1C, 1D). PRMT1 shows both nuclear and cytosolic staining (Figure 1C, 1E) and was also detected at the plasma membrane, mainly in ER-negative tumors (Figure 1E). Moreover, we observed substantial staining of PRMT1 in the stroma of breast tumors as compared to the normal tissue (Figure 1D). Mononuclear cells, fibroblasts and endothelial cells were positively stained for PRMT1 within the stroma (unpublished data).

Altogether, our results indicate that both PRMT1 mRNA and protein levels are higher in breast tumors compared to normal breast tissue, suggesting that PRMT1 could be targeted in BC.

### RNAi-mediated depletion of PRMT1 decreases BC cell viability, clonogenicity and induces DNA damage and apoptosis

To explore the function of PRMT1 in BC cells, we first depleted PRMT1 using two validated siRNAs (PRMT1#7, PRMT1#8) in MDA-MB-468 TNBC cells (Sup. Fig. 2A). We observed that cell viability was significantly decreased upon PRMT1 depletion in MDA-MB-468 cells, in a time-dependent manner (Figure 2A). Similar results were found in other BC cell lines (4 TNBC, 1 Her2+, 2 luminal; Sup.Fig.2B), suggesting that the effect was independent of BC subtype. PRMT1 depletion decreased colony formation in MDA-MB-468 cells under adherent conditions (Figure 2B) or in an anchorage-independent growth assay in soft-agar (Figure 2C), indicating that PRMT1 depletion decreases the tumorigenicity of this TNBC cell line. PRMT1 depletion also decreased colony formation in other BC cells cultured under adherent conditions (Sup. Fig. 2C). Furthermore, we observed a cleavage of caspases 3, 7, and PARP in MDA-MB-468 cells following PRMT1 depletion (Figure 2D), revealing apoptosis induction. This was confirmed in PRMT1-depleted MDA-MB-468 cells using a caspase 3/7 activity assay (Figure 2E) and by extracellular annexin-V staining (Figure 2F). PRMT1 depletion also significantly increased the phosphorylation of histone H2AX (γH2AX), a DNA damage marker (Figure 2D). Induction of apoptosis upon PRMT1 knockdown was confirmed in other BC cell lines (HCC70, MDA-MB-231, SKBr3, T47D; Sup. Fig 2D). Together, these results demonstrate that PRMT1 is required for BC cell survival.

**Figure 2.**
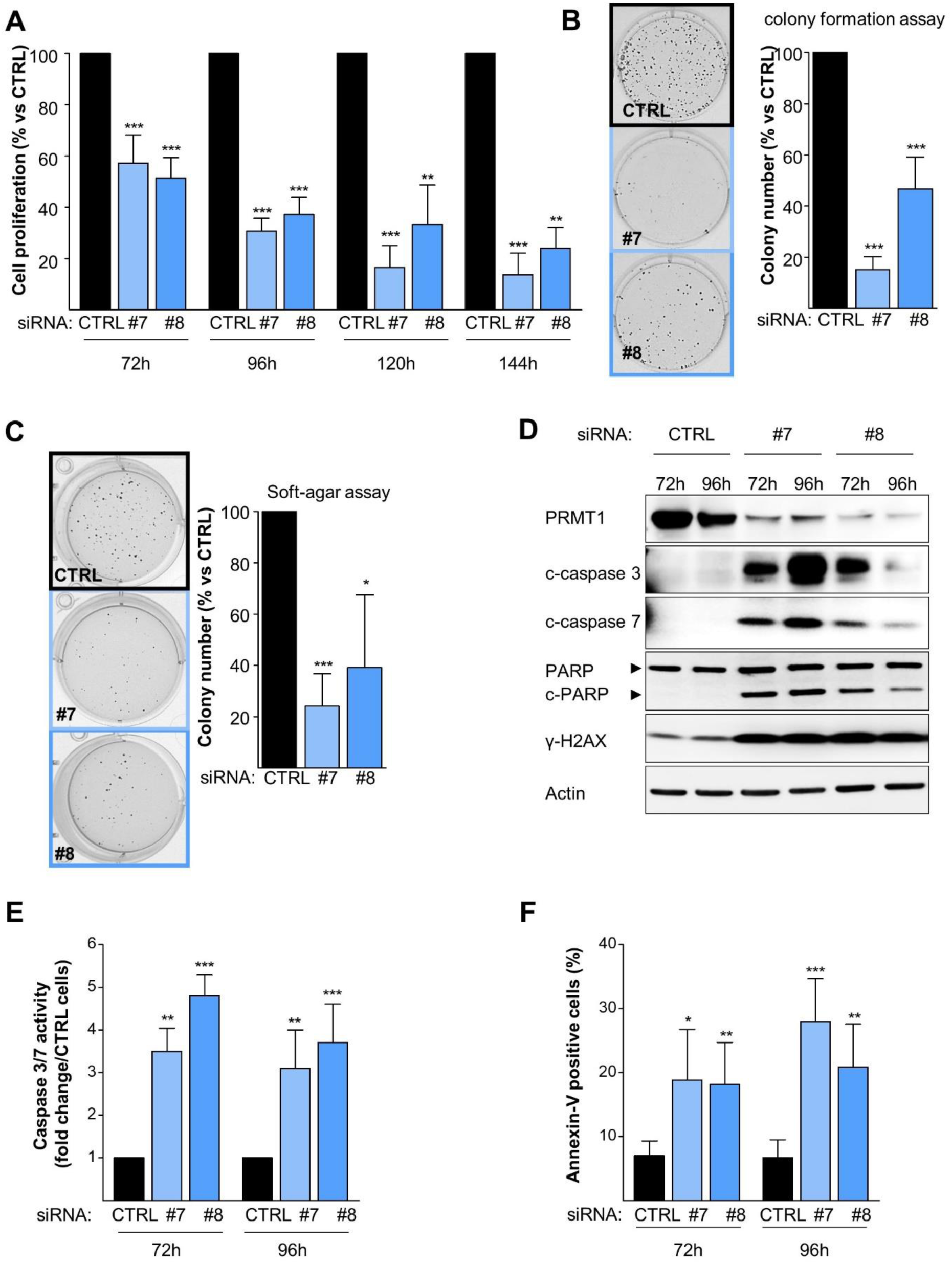
PRMT1 depletion decreases the viability and induces apoptosis of MDA-MB-468 cells. **A,** PRMT1 depletion impairs cell viability (MTT assay). Cells were transfected with control (CTRL, black) or two different PRMT1 siRNAs (#7, #8, blue) for 72h-144h. **B and C,** PRMT1 depletion impairs colony formation when cells are grown on plastic for 13 days (B) or in soft-agar for 4 weeks (C) following siRNA treatment. **D-F**, PRMT1 depletion induces apoptosis. Apoptosis was detected by western blotting using antibodies recognizing the cleaved forms of caspase 7 (c-caspase7), caspase 3 (c-caspase3) and PARP (c-PARP) (D), by caspase 3/7 assay (E) or annexin-V staining (F) after 72h and 96h following siRNA treatment. DNA damage was detected using an anti-γH2AX antibody (D). PRMT1 depletion was verified using an anti-PRMT1 antibody (D). Anti-actin antibody was used as a loading control (D). Results are presented as the percentage (A, B, C, F) or fold change (E) relative to control cells (CTRL). For the quantifications, the data are expressed as the mean ± SD from at least three independent experiments (A, B, C, E, F). Pictures are from a single experiment, representative of three independent experiments (B, C, D). P values are calculated from a Student T test and represented as *p < 0.05; **p < 0.01; ***p < 0.001.

### Type I PRMT inhibitors reduce MDA-MB-468 cell growth

Next, we sought to explore if the enzymatic activity of PRMT1 was necessary for BC cell survival. For this purpose, we used two recently developed type I PRMT inhibitors: MS023 (17) and GSK3368715 (18). Under the tested conditions, both inhibitors decreased the PRMT1-specific histone mark H4R3me2a (Figure 3A) without affecting the methylation of H3R17me2a (by CARM1 and PRMT6) or PABP1 (by CARM1; Sup. Figure 3A). Both inhibitors decreased the viability of MDA-MB-468 cells with similar half-maximal inhibitory concentrations (IC_50_): 2.61±1.75 µM (MS023) and 2.62±1.99 µM (GSK3368715) (Figure 3B). We also observed smaller sized colonies when MDA-MB-468 cells were treated with both inhibitors (Figure 3C).

**Figure 3.**
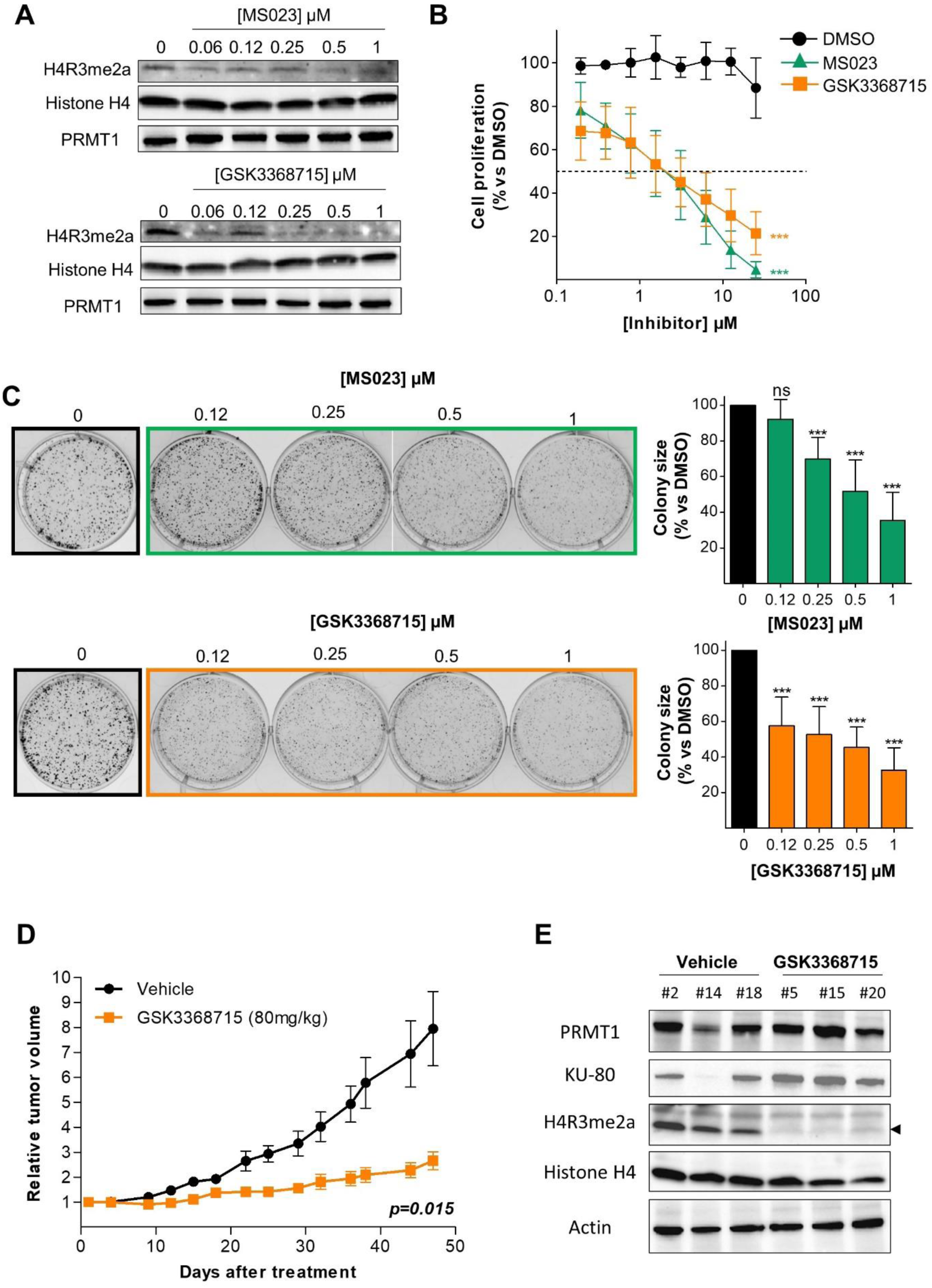
Type I PRMT inhibitors reduce cell viability and tumor growth. **A,** MS023 and GSK3368715 inhibit PRMT1 activity in MDA-MB-468 cells. Cells were treated with MS023 (top) or GSK3368715 (bottom) for 48h and PRMT1 inhibition was assessed by western blotting using anti-H4R3me2a antibody. **B,** Type I PRMT inhibitors decrease MDA-MB-468 cell viability. Cells were treated with DMSO (black circle), MS023 (green triangle) or GSK3368715 (orange square) for 7 days and proliferation was determined by MTT assay. Results are presented as the percentage of cell growth relative to DMSO-treated cells. **C,** Type I PRMT inhibitors reduce the growth of colonies when MDA-MB-468 cells were cultured on plastic for 9 days after MS023 (top) or GSK3368715 (bottom) treatment. Colony size was quantified and is expressed as a percentage relative to DMSO-treated cells (C, right panel). Pictures are from a single experiment representative of three independent experiments (C, left panel). The data are expressed as the mean ± SD from at least three independent experiments (B, C). P values are from a Student t-test and represented as *p < 0.05; **p < 0.01; ***p < 0.001. **D,** GSK3368715 slows tumor growth. Tumors derived from MDA-MB-468 cells were subcutaneously grafted into 12 mice (6 vehicle-treated, black; 6 GSK3368715-treated, orange). Growth curves were obtained by plotting mean relative tumor volume ± SEM as a function of time. P-value was calculated using a Mann-whitney U test. **E,** GSK3368715 inhibits PRMT1 activity, *in vivo*. PRMT1 expression (anti-PRMT1) and activity (anti-H4R3me2a) were analyzed in the tumors excised from 3 vehicle (#2, #14, #18) or GSK3368715 (#5, #15, #20)-treated mice at the end of the experiment (D). Antibodies against histone H4, actin and KU-80 were used as controls.

### Type I PRMT inhibition slows tumor growth in a TNBC xenograft model

We evaluated the anti-tumor effect of inhibiting PRMT1 using GSK3368715, the only type I PRMT inhibitor currently in phase I clinical trial for diffuse large B-cell lymphomas and solid tumors (NCT03666988). To better represent clinical conditions, we engrafted tumors derived from MDA-MB-468 cells into Swiss-nude mice (see materials and methods). GSK3368715 treatment significantly slowed tumor growth (p = 0.015; Figure 3D) with no observed toxicity (Sup. Fig 3B). We confirmed that PRMT1 was indeed inhibited in the tumors at the end of the experiment by observing an increase in pan-monomethylation (Sup. Fig 3C), as reported (34), and a decrease in histone H4R3 methylation (H4R3me2a, Figure 3E).

### PRMT1 regulates the EGFR and Wnt signaling pathways at the transcriptomic level

PRMT1 plays a crucial role in transcriptional regulation (8,11,12). Therefore, we performed transcriptomic analysis of PRMT1 depleted MDA-MB-468 cells to gain insight into the molecular mechanisms that mediate the dependency of BC cells on PRMT1.

MDA-MB-468 cells were transfected with 2 different siRNAs targeting PRMT1 for 24h and 48h and the RNA were analyzed using HTA 2.0 microarrays (Affymetrix). We focused on the genes that were commonly deregulated at 24h and 48h by both siRNAs (Table S2) to perform a gene enrichment pathway analysis using REACTOME (31). The top ranked pathways (according to adjusted p-value) revealed that PRMT1 is involved in several cellular processes including signal transduction pathways, immune system response, lipid metabolism and transcriptional regulation (Figure 4). We focused on EGFR (p = 6.96 x 10^-6^) and Wnt (p = 5.07 x 10^-6^) signaling pathways, which are known to be activated in TNBC (3–5).

**Figure 4.**
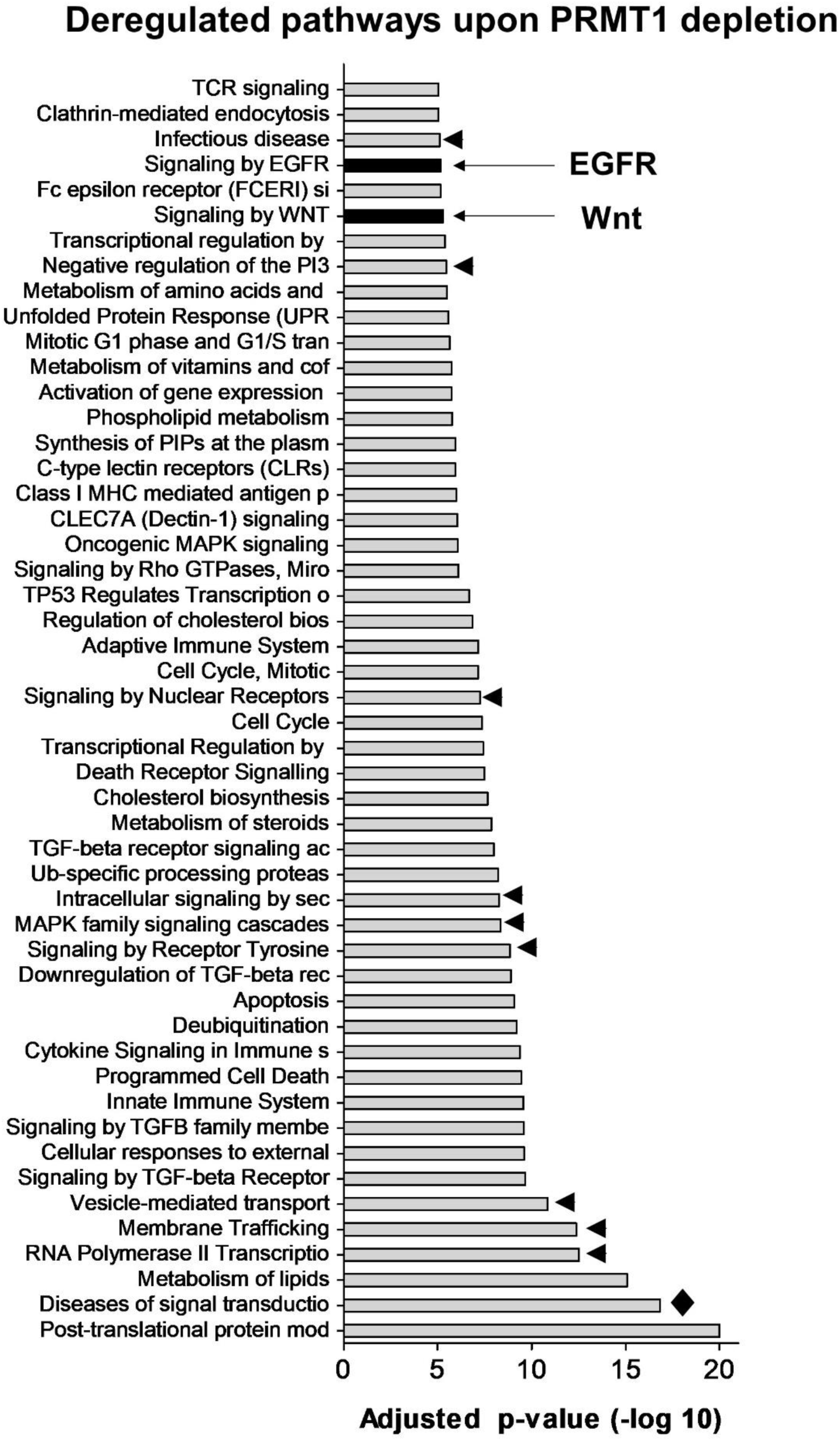
PRMT1 regulates EGFR and Wnt signaling pathways. RNA was extracted from MDA-MB-468 cells transfected with control or PRMT1 (#7, #8) siRNA for 24h and 48h, and analyzed by Affymetrix microarray. Gene enrichment pathway analysis, using the REACTOME database, was performed on the deregulated genes common to both PRMT1 siRNAs. The top 50 deregulated pathways ranked according to their significance (adjusted p-values) is shown. The EGFR and Wnt signalling pathways are highlighted in black. Arrowheads point to pathways including EGFR, and diamond points to pathways including EGFR, LRP5 and PORCN.

We noticed that EGFR itself was less expressed upon PRMT1 depletion in our microarray analysis (Figure 5A) and confirmed this observation by qPCR (Figure 5B). EGFR was also retrieved in several other deregulated pathways (Figure 4, arrowheads and diamond). PRMT1 was directly recruited to two promoter regions of *EGFR* in MDA-MB-468 cells using an anti-PRMT1 antibody (Figure 5C), previously validated for ChIP (35). Further, PRMT1 depletion also decreased the protein expression of EGFR (Figure 5D).

**Figure 5.**
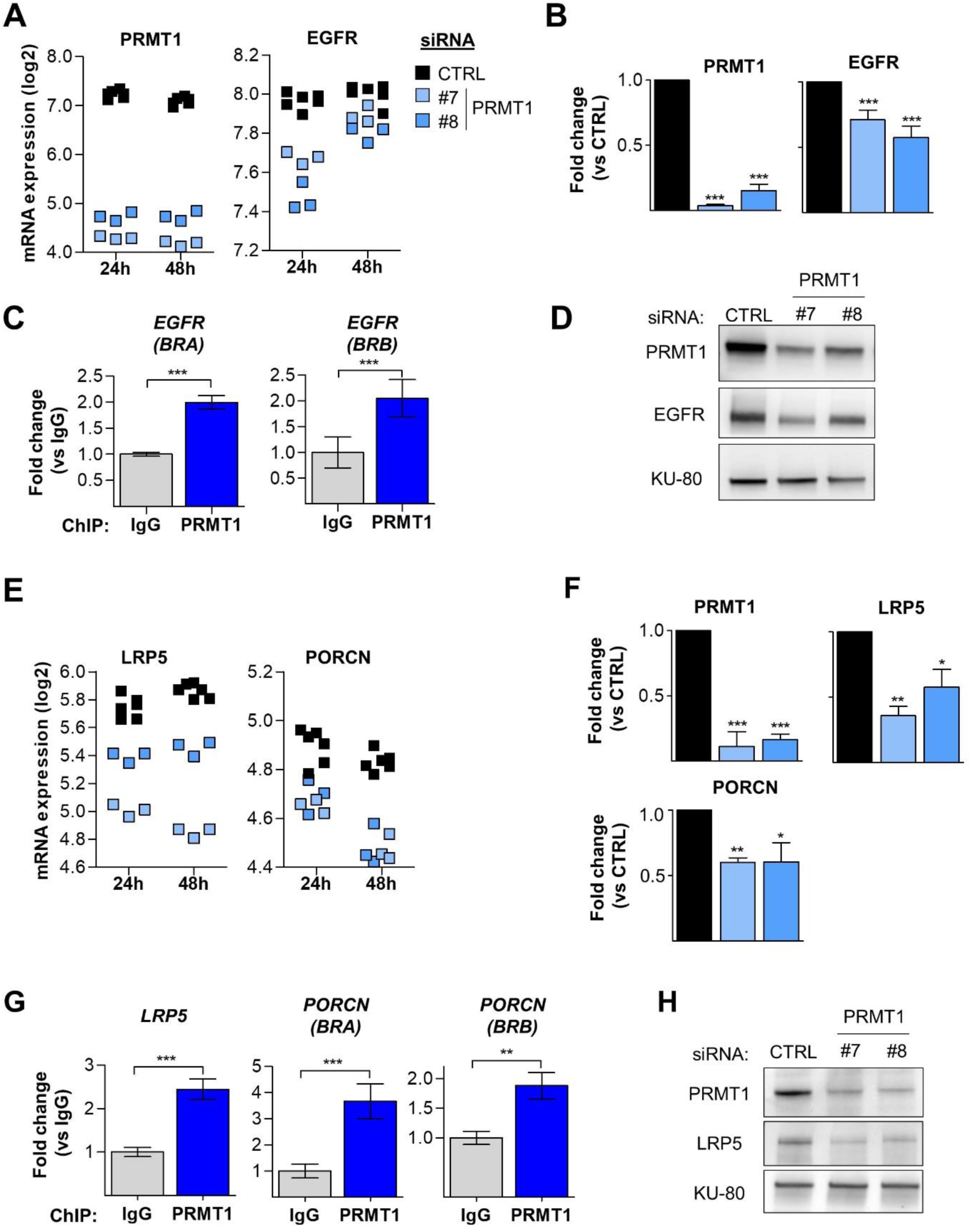
PRMT1 activates the transcription of *EGFR*, *LRP5* and *PORCN* by binding to their promoter. **A and B,** PRMT1 depletion in MDA-MB-468 cells reduces EGFR mRNA expression as shown by Affymetrix microarray (A) and verified by qPCR (B). **C,** PRMT1 binds to two promoter regions (Binding Region A, BRA; BRB) of *EGFR*. **D,** PRMT1 depletion reduces EGFR protein level as shown by western blotting. **E and F,** PRMT1 depletion reduces LRP5 and PORCN mRNA expression as shown by Affymetrix microarray (E) and validated by qPCR (F). **G,** PRMT1 binds to the promoter of *LRP5* and two promoter regions (BRA, BRB) of *PORCN.* **H,** PRMT1 depletion reduces LRP5 protein level as shown by western blotting. MDA-MB-468 cells were transfected with control (black) or two PRMT1 (#7, #8, blue) siRNAs for 24h (A, B, E) and 48h (A, D, E, F, H). mRNA expression was logarithmically transformed (log 2) and each replicate is represented as a single point on the scatter plot (A, E). ChIP were performed using anti-PRMT1 (blue bars) or anti-IgG (grey bars) antibodies using chromatin isolated from MDA-MB-468 cells (C, G). qPCR was performed using primers targeting the promoter regions of *EGFR* (C), *LRP5* (G) and *PORCN* (G). PRMT1 depletion was verified in the Affymetrix microarray (A), by qPCR (B, F) and by western blotting (D, H). Antibody against KU-80 was used as a loading control for the western blots and pictures are representative of at least three independent experiments (D, H). The quantifications are represented as fold change relative to the control siRNA (B, F) or control IgG (C, G) and presented as mean ± SD (B, F) or mean ± SEM (C, G) from three independent experiments. P values from Student t test are represented as *p < 0.05; **p < 0.01; ***p < 0.001.

Our microarray analysis revealed two key players of the Wnt signaling pathway, LRP5 and Porcupine (PORCN), to be less expressed following PRMT1 depletion (Figure 5E). LRP5 and PORCN were also found in the second-top deregulated pathway (Figure 4, diamond). We validated this by qPCR (Figure 5F) and identified by ChIP analysis that PRMT1 binds to the promoter of *LRP5* and two regions of the *PORCN* promoter (Figure 5G). The expression of LRP5 was also decreased at the protein level after PRMT1 depletion (Figure 5H). We could not assess the protein expression of porcupine due to the lack of suitable antibodies for western blotting.

Overall, these results indicate that PRMT1 regulates the expression of EGFR, LRP5 and PORCN by directly binding to their promoter regions.

### PRMT1 activates the canonical Wnt signaling pathway

We hypothesized that PRMT1 could be an activator for the Wnt pathway as both LRP5 and PORCN are required for its activation. We first assessed Wnt activity by analyzing the expression of the three Wnt target genes (*AXIN2*, *APCDD1*, and *NKD1*) that are the most upregulated in Wnt3a-stimulated MDA-MB-468 cells (30). We observed that PRMT1 depletion reduced the expression of these three Wnt target genes (Figure 6A, top panel). By using the gold standard β-catenin activated reporter (BAR) assay, we confirmed that PRMT1 depletion decreased Wnt signaling activity (Figure 6B). siRNA targeting LRP6 was used as a positive control in both assays (Figure 6A, 6B).

**Figure 6.**
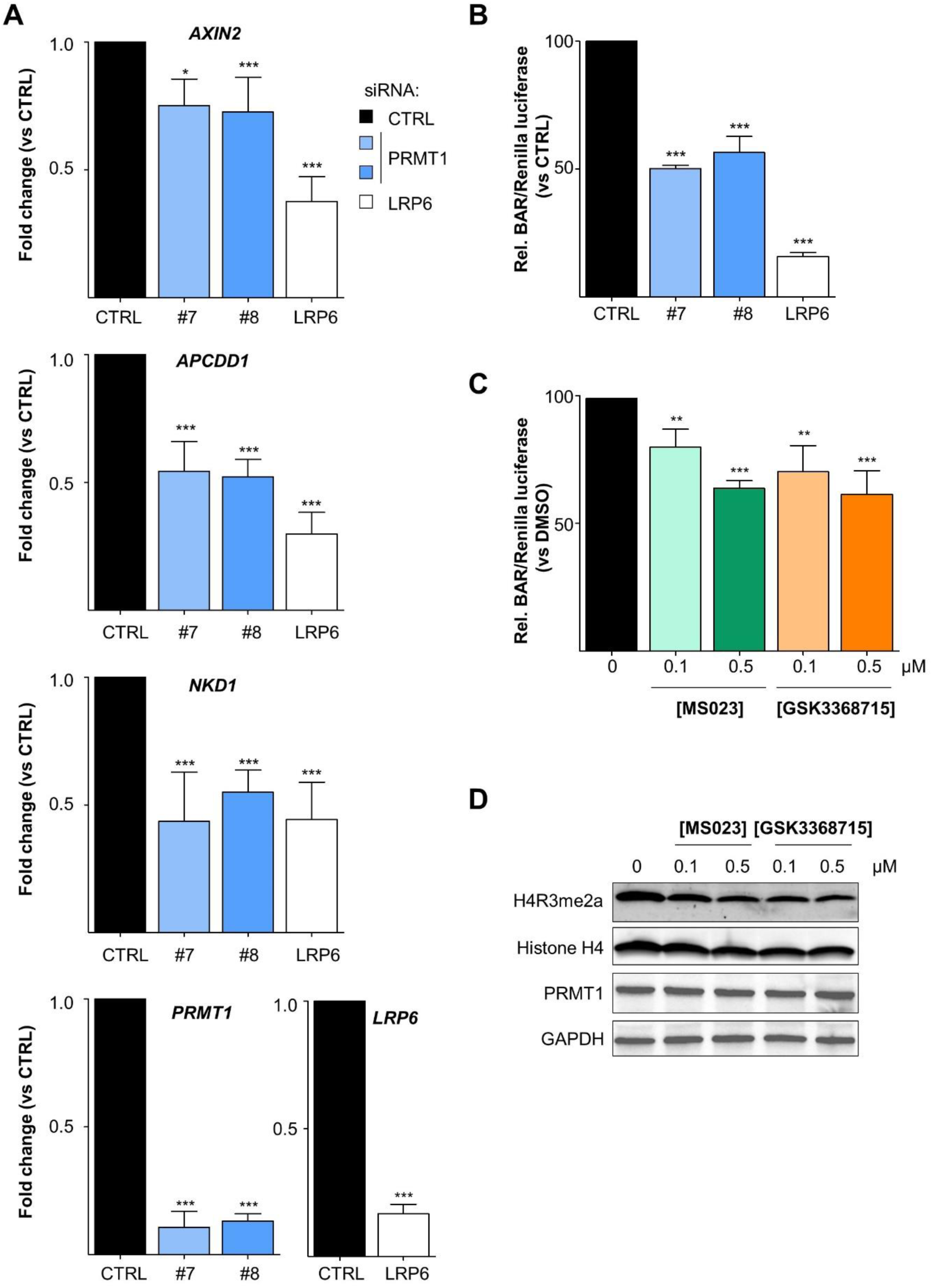
PRMT1 activates the canonical Wnt signaling pathway. **A and B,** PRMT1 depletion decreases Wnt signalling actvity. MDA-MB-468 cells were transfected with control (CTRL, black), two PRMT1 (#7, #8, blue) or LRP6 (white) siRNA for 48h (A, B), and then co-transfected with plasmids coding for BAR-firefly luciferase and Renilla luciferase for 24h (B), before Wnt3a stimulation for 6h (A, B). The expression of *AXIN2*, *APCDD1*, *NKD1* (Wnt target genes), *PRMT1* and *LRP6* were quantified by qPCR (normalized to actin) (A). The relative luciferase signal (firefly luciferase/ Renilla luciferase) is represented as a percentage normalised to the control siRNA (CTRL) (B). siRNA targeting LRP6 was used as a positive control (A, B). **C,** Type I PRMT inhibitors decrease Wnt signaling activity. MDA-MB-468 cells were treated with 0.1μM or 0.5μM of MS023 (green) or GSK3368715 (orange) for 48h, and then co-transfected with plasmids coding for BAR-firefly luciferase and Renilla luciferase for 24h, before Wnt3a stimulation for 6h. The relative luciferase signal (firefly luciferase/ Renilla luciferase) is represented as a percentage normalised to the DMSO treated cells (black). **D,** PRMT1 inhibition was verified in this experiment (C) by western blotting using anti-H4R3me2a antibody. Anti-histone H4, PRMT1, and GAPDH were used as loading controls. All quantifications are represented as fold change (A) or percentage (B, C) relative to the control. The data are expressed as the mean ± SD from at least three independent experiments (A, B, C). P values from Student t-test are represented as *p < 0.05; **p < 0.01; ***p < 0.001.

Next, we checked if PRMT1 enzymatic activity was involved in the regulation of Wnt pathway. MDA-MB-468 cells were treated for 3 days with low doses of MS023 or GSK3368715 (0.1μM and 0.5μM) and then stimulated for 6h with Wnt3a, before assessing Wnt activity using the BAR assay (Figure 6C). Both type I PRMT inhibitors decreased Wnt activity in a dose-dependent manner (Figure 6C). PRMT1 was inhibited under these conditions (Figure 6D).

Collectively, this demonstrates that PRMT1 and its activity are involved in the activation of the canonical Wnt pathway in MDA-MB-468 cells.

### Type I PRMT inhibitors show synergistic interactions with erlotinib or chemotherapies

The rationale of drug combinations is to improve efficacy, limit side-effects and reduce the risk of drug resistance. First, we combined the two type I PRMT inhibitors with chemotherapies (cisplatin, camptothecin, cyclophosphamide, taxanes) used in the clinic to treat TNBC patients. MDA-MB-468 cells were treated with varying concentrations of the drugs, starting from about 2 x IC_50_ (Table S1) for 7 days (equivalent to 4 mitotic cycles) and cell viability was assessed using CellTiterGlo assay. We applied the Loewe additivity model using the Combenefit software (36) to determine the nature (synergy/additivity/antagonism) of the drug interactions. We used this model as it allows the possibility to analyze two drugs that may act on same pathways (37). Both type I PRMT inhibitors synergized with cisplatin (Figure 7A, Sup. Fig 4A), camptothecin (Figure 7B, Sup. Fig 4B) and cyclophosphamide (Figure 7C, Sup. Fig 4C), but not with docetaxel (Sup. Fig 5A) or paclitaxel (Sup. Fig 5B).

**Figure 7.**
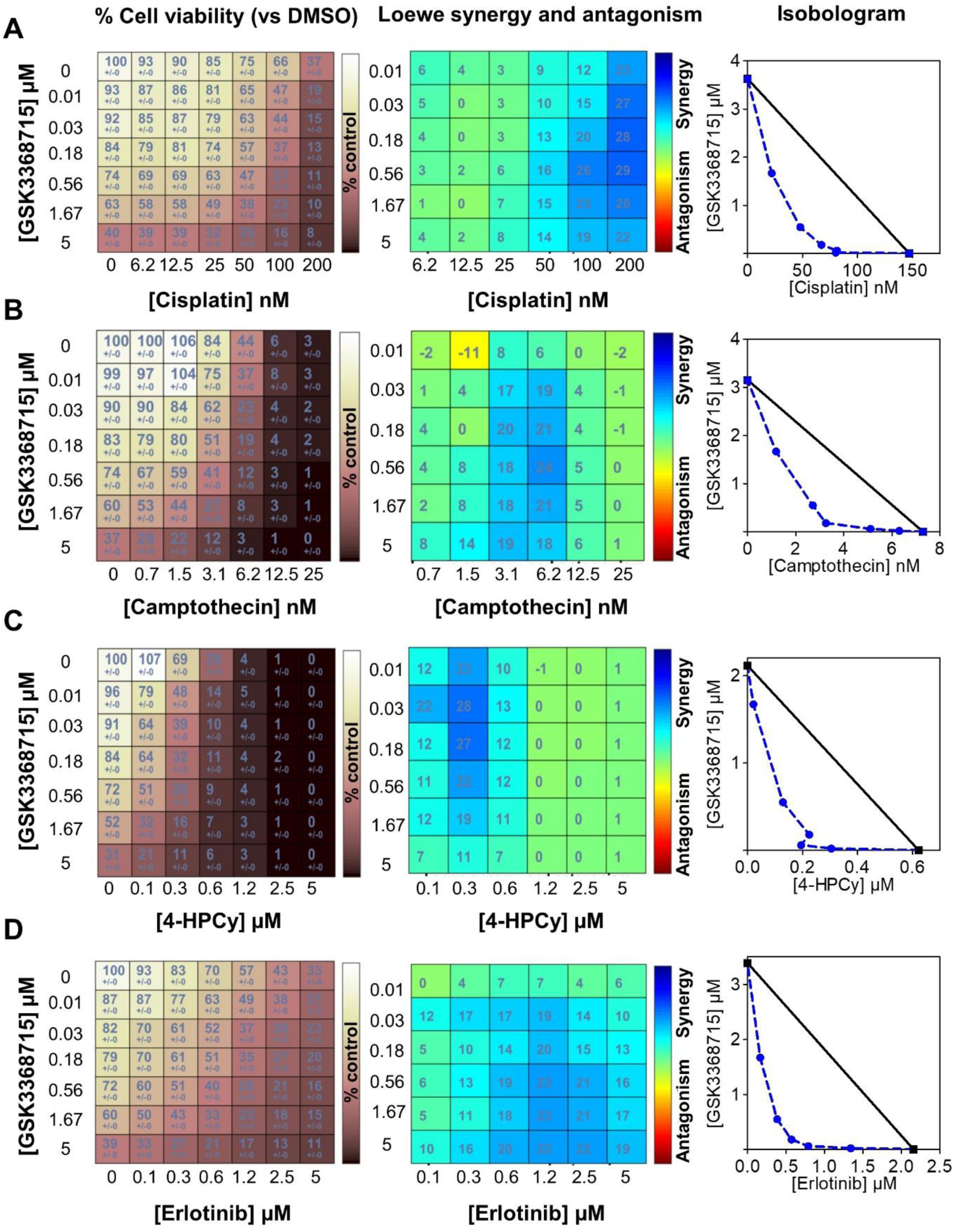
Synergistic interactions between Type I PRMT inhibitor (GSK3368715) and chemotherapies (A, B, C) or erlotinib (D). MDA-MB-468 cells were seeded in 96-well plates, treated with the indicated drugs for 7 days (equivalent to 4 doubling times) and cell viability was measured by CellTiterGlo assay. GSK3368715 was serially diluted three-fold and cisplatin (A), camptothecin (B), 4-hydroperoxy cyclophosphamide (4-HPCy; C), erlotinib (D) were serially diluted two-fold (concentrations indicated in the figure). The drug interactions were calculated using the Loewe model on the Combenefit software. Cell viability (% compared to DMSO-treated cells, left panel), synergy matrix as calculated using the Loewe excess model (middle panel), and isobolograms (right panel) for each drug pair are indicated. Presented data are representative of three independent experiments.

As EGFR is highly expressed in TNBC (3), we also evaluated the potential of combining type I PRMT inhibitors with an EGFR inhibitor (erlotinib) and observed a high synergy (Figure 7D, Sup. Fig 4D). These combinations may represent promising alternative therapeutic strategies for TNBC patients.

## Discussion

The efficacy of breast cancer therapies has considerably improved; however, TNBC maintain a poor prognosis compared with other subtypes and are typically correlated with increased recurrence and worse survival. Finding alternative treatments to chemotherapy remains a priority to treat TNBC patients to avoid relapses.

PRMTs are overexpressed in various cancer types and are emerging as attractive therapeutic targets (8–13). Consequently, several PRMT inhibitors have been developed and some PRMT5 and type I PRMT inhibitors are being evaluated in clinical trials (10).

At the RNA level, we found that PRMT1 is more expressed in BC when compared to the normal breast tissue, aligning with previous studies which did not consider BC heterogeneity (38, 39). PRMT1 mRNA correlates positively with Ki-67. Consequently, the highest PRMT1 expression was found in TNBC, the most proliferative BC subtype, and this could be a result of DNA copy number gain. High PRMT1 mRNA expression correlates with poor prognosis in all breast tumors, as reported (33, 40), as well as within LA and LB subtypes. In contrast, TNBC expressing the highest level of PRMT1 (most proliferative) display better RFS, most likely because they respond better to chemotherapy.

At the protein level, PRMT1 is more expressed in BC compared to normal tissue, confirming previous reports (33, 40). Here, we accounted for BC heterogeneity and found that PRMT1 protein is expressed at similar levels in the different BC subtypes. We observed both nuclear and cytosolic staining for PRMT1 which is in apparent contrast to a study showing mainly cytosolic localization (40), using an antibody that also recognizes the C-terminus of PRMT1, therefore detecting all its isoforms (41). Several PRMT1 splice variants have been described which show cytoplasmic and/or nuclear localization (39), therefore it may not be surprising to detect PRMT1 in both compartments. We also detected PRMT1 at the plasma membrane, preferentially in the ER-negative BC subtypes, possibly since it interacts with some transmembrane receptors such as EGFR (19, 20) or IGF-1R (42). However, we cannot exclude that the PRMT1 antibody we used, recognizes the plasma membrane associated-PRMT8, although it is brain-specific, as it shares 80% homology with PRMT1.

Transcriptomic analysis highlighted several pathways regulated by PRMT1. Here, we focused on two pathways that are known to be activated in TNBC (3–5). PRMT1 has been previously observed to modulate EGFR signaling by two mechanisms: (i) by methylating histone H4 (H4R3me2a) on its promoter in colorectal cancer (CRC) (22) and glioblastoma cells (23) and (ii) by methylating EGFR in CRC and TNBC cells (19, 20). Here, we demonstrate that PRMT1 activates the transcription of EGFR by directly binding to its promoter.

The role of PRMT1 on Wnt signaling is ambiguous since PRMT1 can be both an activator or an inhibitor of this pathway. On one hand, PRMT1 can inhibit Wnt signaling by methylating two antagonists (i) Axin (in HEK293 and L929 cell lines) (26) and (ii) Dishevelled (in HEK293, B2b, and F9 cell lines) (27). On the other hand, PRMT1 can activate the Wnt signaling pathway by methylating two Dishevelled-binding components: G3BP1 (in F9 cells) (24) and G3BP2 (in F9, HEK293 and SW380 cells) (25). Therefore, the role of PRMT1 on Wnt signaling may be context dependent. Here, we show that PRMT1 regulates the Wnt signaling pathway at the transcriptomic level. Indeed, we found that PRMT1 activates the transcription of two main components of the Wnt pathway: *LRP5* and *PORCN*, by binding to their promoter. Furthermore, we demonstrate that PRMT1 activates the canonical Wnt signaling pathway. Additionally, PRMT1 enzymatic activity could be required as type I PRMT inhibitors reduce Wnt signaling pathway. Hence, PRMT1 could activate the pathway by directly methylating Wnt components or methylating histones on their promoters. Together, this implies that PRMT1 may regulate the Wnt signaling pathway by regulating the amounts of LRP5 available at the plasma membrane, and by controlling the Porcupine-dependent post-translational modification of Wnt ligands, which is required for their secretion.

As PRMT1 is highly expressed in BC, we evaluated its potential as a therapeutic target. We found that PRMT1 depletion (i) decreased BC cell viability, (ii) blocked their clonogenic potential, and (iii) induced DNA damage and apoptosis in various cell lines of different BC subtypes. This is in accordance with previous reports in TNBC (20,33,43,44) and luminal (35,44,45) BC cell lines as well as cell lines of other cancer types (22,46–49). We next addressed the question whether the enzymatic activity of PRMT1 was required for BC cell survival. To date, there are no PRMT1 specific small-molecule inhibitors, but rather inhibitors that target all type I PRMTs, with some selectivity towards PRMT1, PRMT6, and PRMT8 (17, 18). GSK3368715 targets these three PRMTs at similar IC_50_ (18) whereas PRMT6 and PRMT8 are more sensitive than PRMT1 to MS023 (17). We found that both inhibitors decrease cell viability and the growth of colonies in MDA-MB-468 cells aligning with previous studies in other cancer cell lines (18,50–52). Together, we found that PRMT1 and its enzymatic activity are required for BC cell survival; however, we cannot rule out the influence of PRMT6 activity when using these inhibitors in our BC cell lines.

When assessing these inhibitors in combination with chemotherapies used in the clinic to treat TNBC patients, we observed synergistic interactions with cisplatin, cyclophosphamide, and camptothecin, but not with docetaxel and paclitaxel in MDA-MB-468 cells. Notably, these synergistic interactions occurred at doses lower than the IC_50_ of each drug, therefore potentially minimizing their cytotoxic side-effects when used in combination *in vivo*. MS023 treatment was shown to sensitize ovarian cancer cells to cisplatin (53), CRC cells to SN-38, a camptothecin derivative (54). In order to generalize our findings, these combinations must be tested in additional TNBC cell lines.

The highest synergy was observed when we combined both type I PRMT inhibitors with erlotinib in MDA-MB-468 cells, a cell line overexpressing EGFR. It would be valuable to test this combination in other TNBC cell lines to verify whether this synergy is associated with EGFR overexpression. Indeed, we have previously reported a synergistic interaction between erlotinib and a PRMT5 inhibitor, independently of the EGFR expression status of TNBC cell lines (16). Although EGFR is overexpressed in TNBC, targeting EGFR on its own has shown only a modest effect in clinical trials in TNBC patients (3). Considering our results, it may be beneficial to combine EGFR and PRMT inhibitors to treat TNBC. However, this hypothesis must be tested *in vivo* in various TNBC patient-derived xenograft (PDX) models. Additional studies have reported that type I PRMT inhibitors synergize with inhibitors targeting PARP in lung cancer (55), PRMT5 in leukemia, pancreatic and lung cancer (18,56,57), FLT3 kinase in leukemia (50, 58) or anti-PD-1/PD-L1 in various cancer types (59, 60). Altogether, this also highlights the potential clinical relevance of combining type I PRMT inhibitors with targeted-therapies.

We performed pre-clinical studies to explore the translational relevance of targeting PRMT1 using GSK3368715, which is being evaluated in a phase I clinical trial. We show that this inhibitor significantly reduced tumor growth, in accordance with a previous study (18). However, there are two major differences between the two studies: (i) we engrafted tumors derived from MDA-MB-468 cells in contrast to Fedoriw et al., who directly injected these cells into the mice, (ii) we used a lower inhibitor dose (80mg/kg) as compared to them (150mg/kg) (18). Type I PRMT inhibitors have also been shown to decrease tumor growth in other cancer types such as lymphoma (18), pancreatic (18, 34), hepatocellular carcinoma (52) and colon (51, 59) cancer. Therefore, targeting type I PRMTs could represent a new treatment strategy in various cancer types, including BC. Additionally, we have evidence supporting the idea that combining the type I PRMT inhibitors with chemotherapies or targeted therapies, could be beneficial for the treatment of TNBC. This must be evaluated in various TNBC PDX models to account for the inter- and intra-tumor heterogeneity observed within TNBC (2). Intra-tumor heterogeneity poses a major challenge in treating TNBC patients because of a subpopulation of cells resistant to chemotherapy leading to residual disease and relapse (2). These chemo-resistant cells are believed to be fueled by developmental pathways such as the Wnt signaling pathway (2,4,5), hence, inhibiting PRMT1 may eradicate these resistant cells. Therefore, addressing whether the drug combinations identified here (*in vitro*) could overcome relapse in chemo-resistant TNBC PDX models would be clinically valuable.

## Acknowledgments

We are grateful to Bérengère Marty-Prouvost for her experimental contribution, particularly for the sample preparation used for the transcriptomic analysis. We thank Virginie Maire for helpful advice, and Dr. Faisal Mahmood and Dr. Ramon Garcia-Areas for enriching discussion regarding the Wnt signaling pathway. We thank Ausra Surmieliova for performing the PRMT1-ChIP experiments, and Laetitia Lesage for scanning the PRMT1 IHC slides.

