## Supplementary figures and methods for "PRMT1 regulates EGFR and Wnt signaling pathways and is a promising target for combinatorial treatment of breast cancer"

[1]Institut Curie, PSL Research University, Paris, France; [2]Translational Research Department; [3]Breast Cancer Biology Group; [4]Pre-clinical investigation laboratory; [5]University of Lyon, Inserm U1052, CNRS UMR5286, Cancer Research Center of Lyon, Lyon, France; [6]Institut Curie Hospital, Department of Diagnostic and Theranostic Medicine, Platform of Experimental Pathology, Paris, France; [7]Genomics Core Facility.

#### Supplementary figures and legends

#### Supplementary Figure S1.

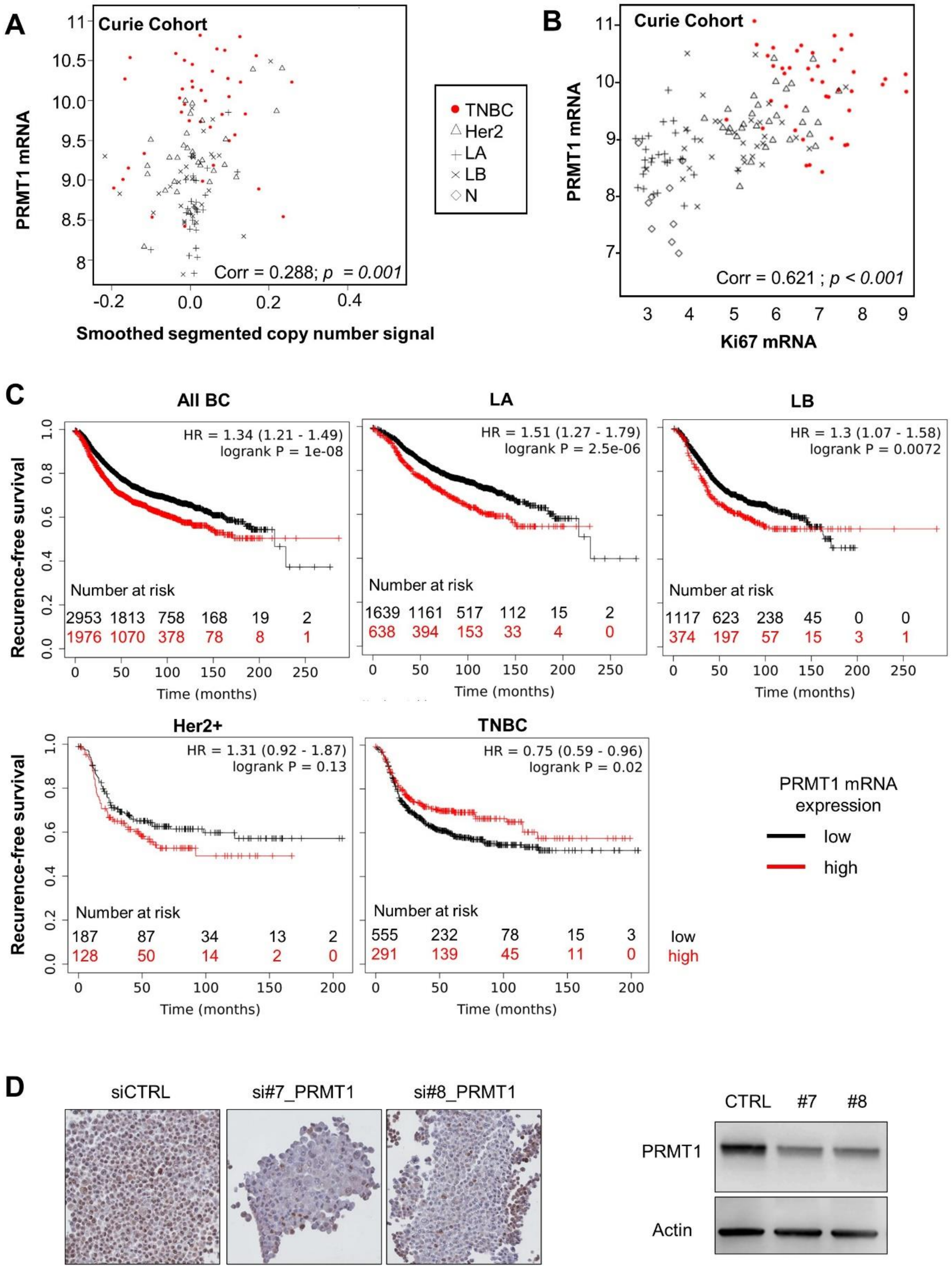


19

**Supplementary Figure S1. Correlation and survival analyses and validation of PRMT1 antibody for IHC. A,** PRMT1 mRNA expression correlates with DNA copy number in the whole population of breast cancer in the Curie cohort (Spearman correlation). **B,** PRMT1 and Ki67 (proliferative marker) mRNA expression positively correlate in the whole breast cancer population of the Curie cohort (Spearman correlation). **C,** PRMT1 mRNA expression correlates with prognosis in BC. Recurrence-free survival based on PRMT1 mRNA expression (Affy probe ID: 206445_s_at) was obtained from the Kaplan-Meier (KM) plotter website (http://kmplot.com). Best performing cutoff option was used: all BC (n=4929), Luminal B (LB, n=1491), Luminal A (LA, n=2277), Basal for TNBC (TN, n=846), and Her2+ (n=315). Hazard ratio with 95% confidence interval and log-rank P values were calculated and significance threshold was set at $p < 0.05$. Of note, a similar figure plotting PRMT1 expression (median cutoff) vs RFS in the whole BC population has been previously published (1) but with a lower number of samples (4929 in our study compared to 3951 in their article). **D,** Validation of PRMT1 antibody for IHC staining. **Left panel,** PRMT1 antibody used for IHC (Fig. 1C) was validated using AFA-fixed cell pellets from MDA-MB-468 cells treated with control siRNA (CTRL) or two siRNAs targeting PRMT1 for 72 h (#7, #8). **Right panel,** PRMT1 depletion was verified by western-blotting using anti-PRMT1 antibodies. Anti-actin antibody was used as a loading control.

### Supplementary Figure S2.

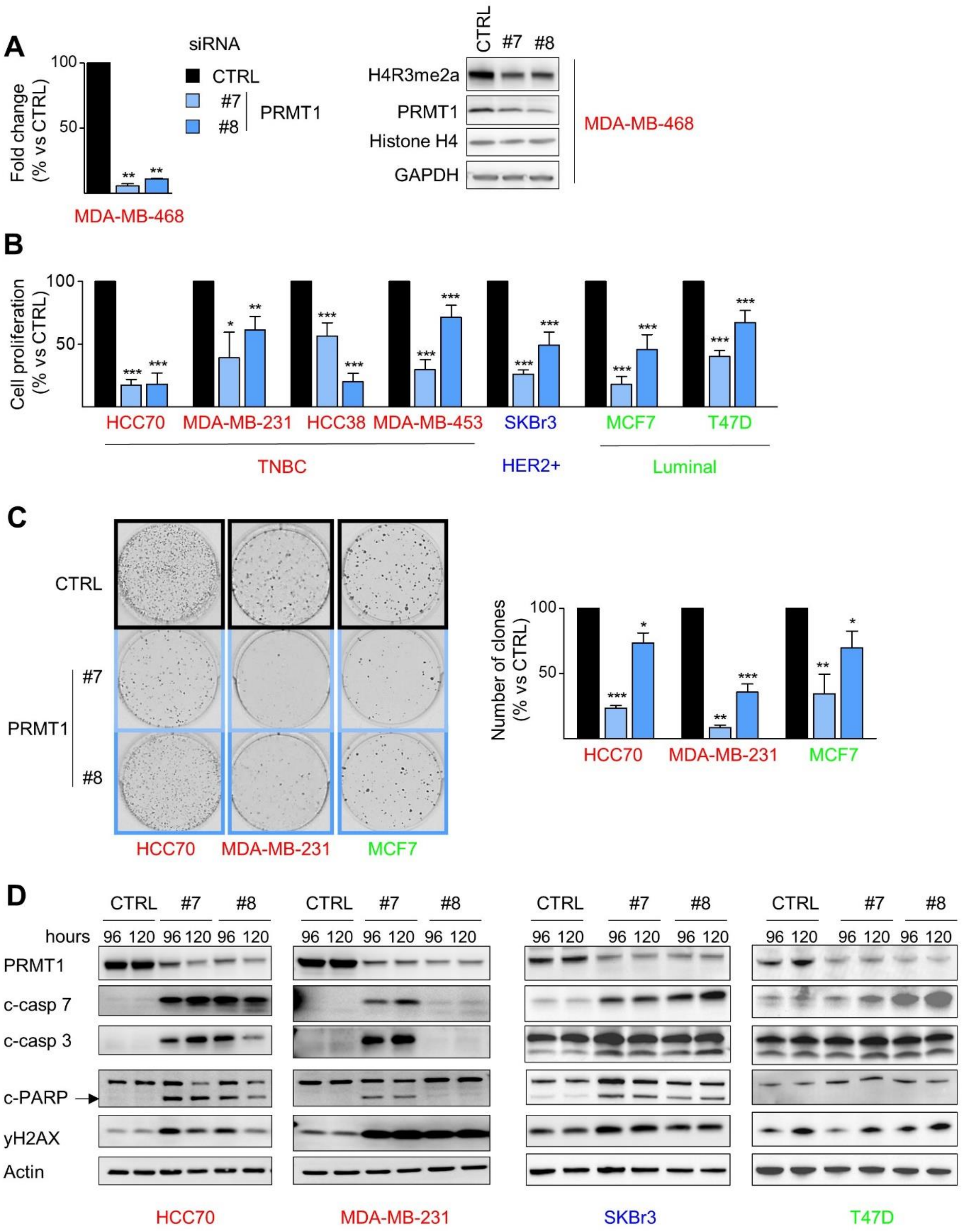


**Supplementary Figure S2. PRMT1 depletion decreases cell viability, colony forming ability and induces apoptosis in various breast cancer cell lines. A,**

Validation of PRMT1 siRNAs. MDA-MB-468 cells were treated with control (CTRL, black) or two PRMT1 (#7, #8, blue) siRNA for 48h. PRMT1 expression was detected at the mRNA (by RTqPCR, normalization by actin; left panel) and protein (by western blotting, right panel) levels. The methylation of histone H4 on Arg 3 (H4R3me2a) was used to measure PRMT1 activity and anti-histone H4 and anti-GAPDH antibodies were used as loading controls. **B,** PRMT1 depletion decreases the viability of breast cancer cells. TNBC (red), Her2+ (blue), and luminal (green) cells were transfected with control (CTRL, black) or two PRMT1 (#7, #8, blue) siRNA for 144h and cell viability was measured by an MTT or WST1 assay. **C,** PRMT1 depletion decreases colony formation. TNBC (red) and luminal (green) cells were transfected with control (CTRL, black) or two PRMT1 (#7, #8, blue) siRNAs, and then cultured on plastic for 6 mitotic cycles equivalent to 14 (HCC70), 7 (MDA-MB-231) or 12 (MCF7) days. A representative image (left panel) and the quantifications (right panel) are shown. **D,** PRMT1 depletion induces apoptosis in BC cells. TNBC (red), Her2+ (blue) and luminal (green) cells were transfected with control (CTRL) or two PRMT1 (#7, #8) siRNA for 96h or 120h. Apoptosis was detected by western blotting using antibodies recognizing the cleaved forms of caspase 7 (c-casp7), caspase 3 (c-casp3) and PARP (c-PARP). DNA damage was detected using an anti-γH2AX antibody. Anti-actin antibody was used as a loading control. Arrow indicates the cleaved form of PARP, while the upper band corresponds to total PARP protein. Results are presented as the percentage (B, C) or percent fold change (A) relative to control cells (CTRL). All data are expressed as the mean ± SD from at least three independent experiments (A, B, C). Pictures are from a single experiment, representative of three independent experiments (A, C, D). P values from a Student t test are represented as $*p < 0.05$; $**p < 0.01$; $***p < 0.001$.

#### Supplementary Figure S3.

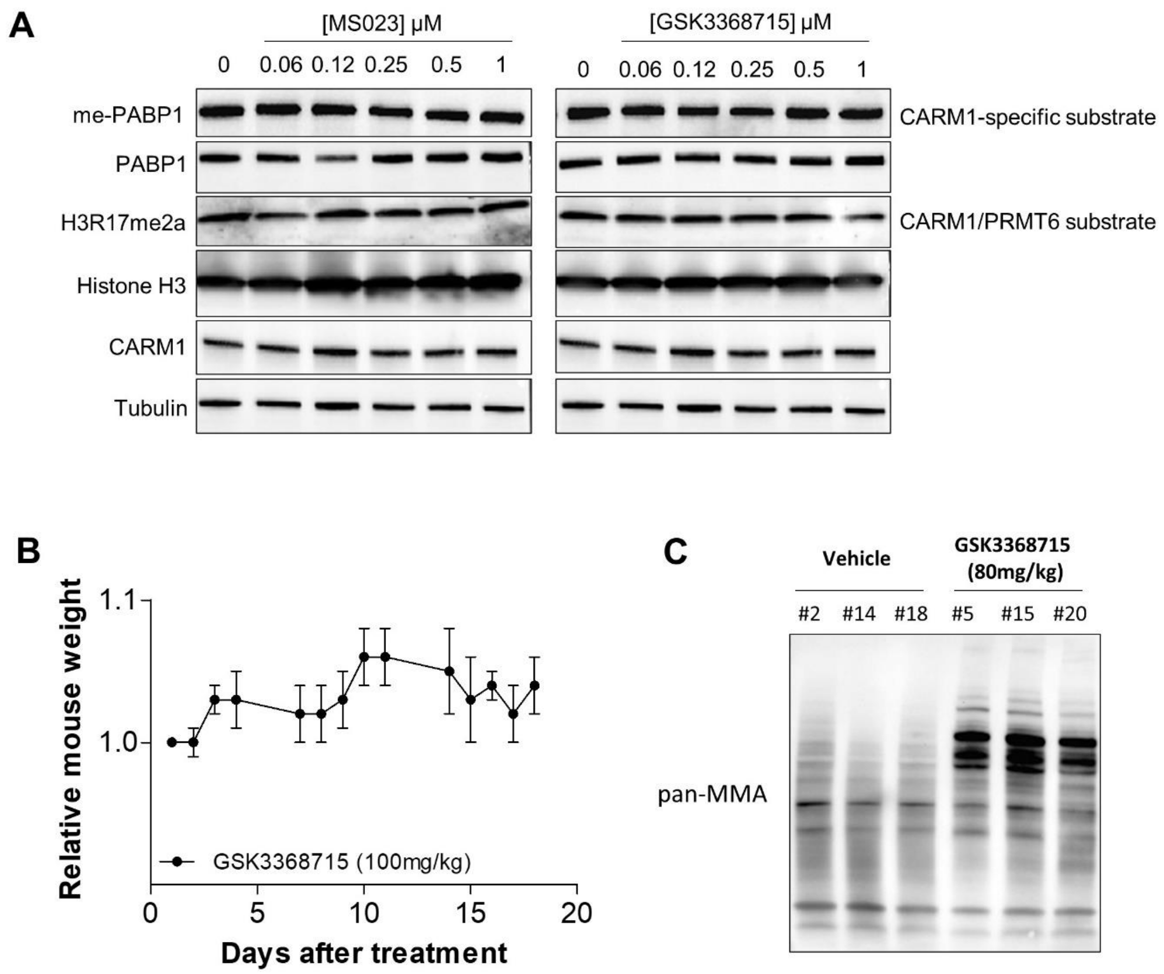


**Supplementary Figure S3. GSK3368715 treatment shows no toxicity and increases global monomethylation in mice. A,** Type I PRMT inhibitors do not affect CARM1 and PRMT6 activity under the tested conditions. MDA-MB-468 cells were treated with varying concentrations of MS023 or GSK3368715 for 48h. Methylation of PABP1 (me-PABP1) was used to measure CARM1 activity and histone H3 methylation (H3R17me2a) to assess CARM1 and PRMT6 activity. Anti-PABP1, histone H3, CARM1, and tubulin antibodies were used as loading controls. This figure is part of Figure 3A. **B,** GSK3368715 treatment is not toxic for mice at the tested dose.

GSK3368715 was administered in Swiss-nude mice (n=3) at 100 mg/kg per-os, once daily for 18 days. Treatment was not associated with any mortality or body weight loss during this period. **C,** GSK3368715 treatment (80mg/kg) increases global monomethylation, *in vivo*. Total monomethylation was detected by western blotting using anti-pan monomethylated (pan-MMA) antibodies in the tumors excised from 3 vehicle (#2, #14, #18)- or GSK3368715 (#5, #15, #20)-treated mice at the end of the experiment.

#### Supplementary Figure S4.

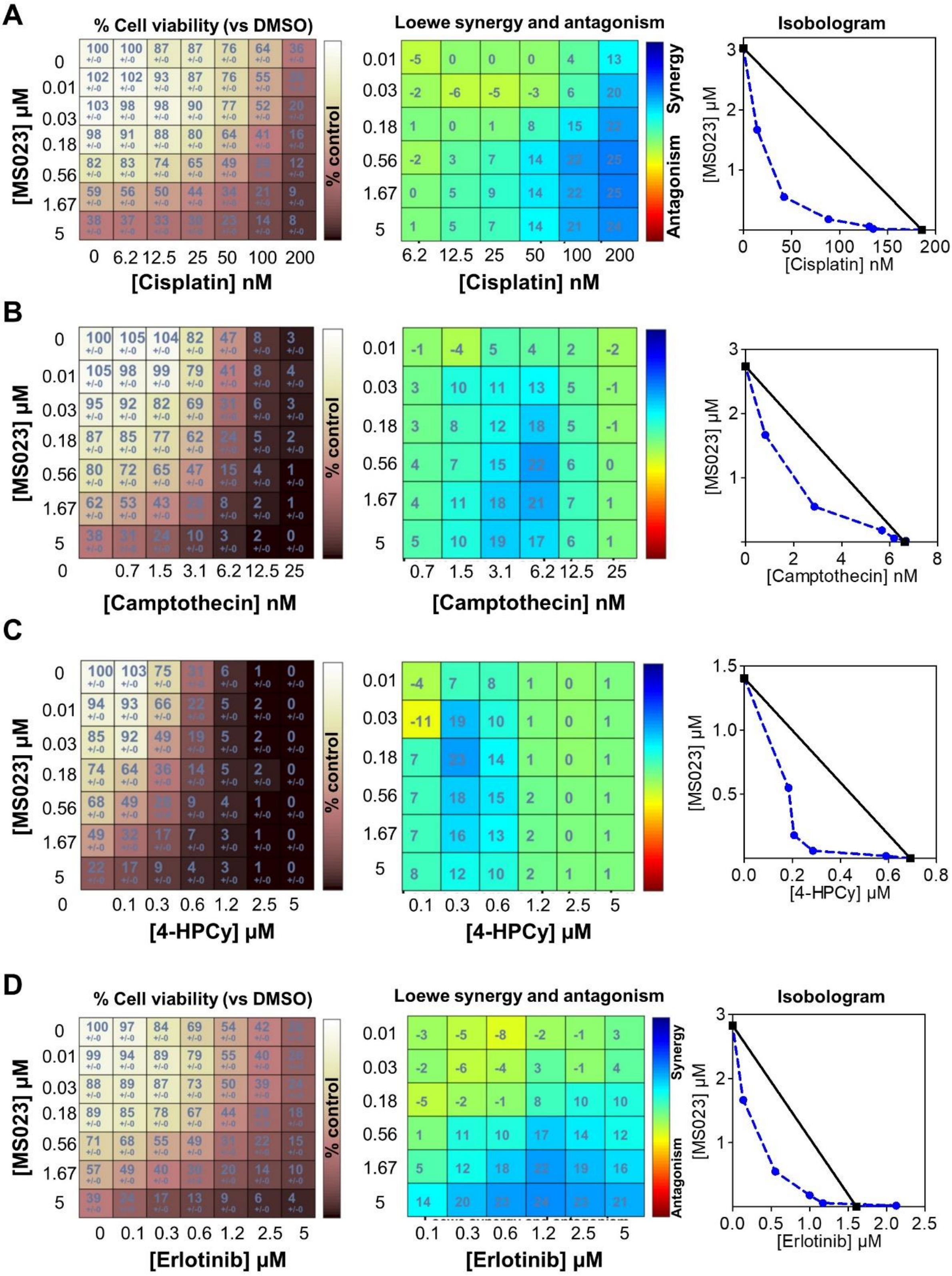

**Supplementary Figure S4. Synergistic interactions between Type I PRMT inhibitor (MS023) and chemotherapies (A, B, C) or erlotinib (D).** MDA-MB-468 cells were seeded in 96-well plates, treated with the indicated drugs for 7 days (equivalent to 4 doubling times), and cell viability was measured by CellTiterglo assay. MS023 was serially diluted three-fold and cisplatin (A), camptothecin (B), 4-hydroperoxy cyclophosphamide (4-HPCy; C), erlotinib (D) were serially diluted two-fold (concentrations indicated in the figure). The drug interactions were calculated using the Loewe model on the Combenefit software. Cell viability (% compared to DMSO-treated cells, left panel), synergy matrix as calculated using the Loewe excess model (middle panel), and isobolograms (right panel) for each drug pair are indicated. Presented data are representative of three independent experiments.

#### Supplementary Figure S5.

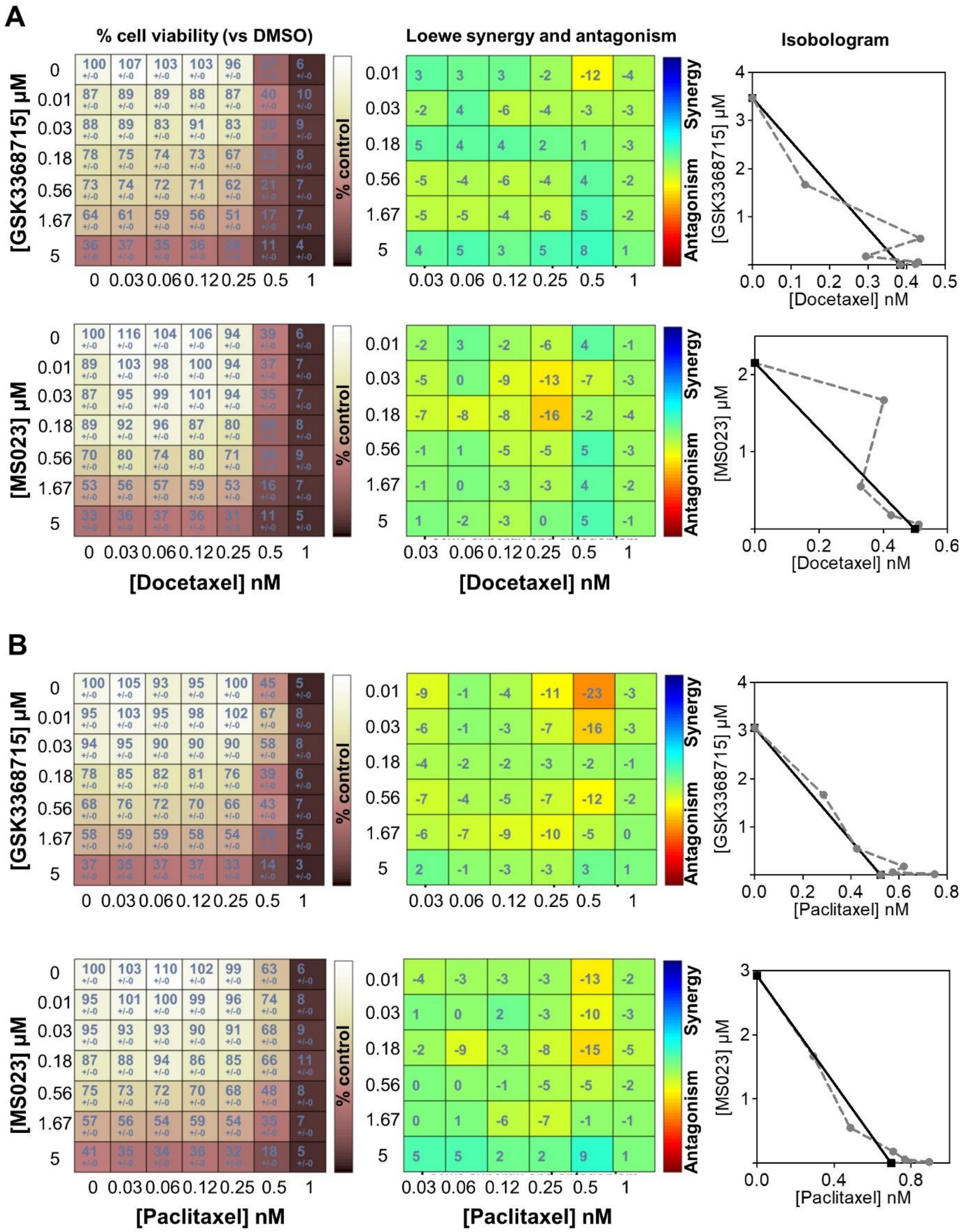

**Supplementary Figure S5. Additive interactions between Type I PRMT inhibitors and taxanes. A and B,** MDA-MB-468 cells were seeded in 96-well plates, treated with the indicated drugs for 7 days (equivalent to 4 doubling times) and cell viability was measured by CellTiterglo assay. Type I PRMT inhibitors (MS023, GSK3368715) were serially diluted three-fold and docetaxel (A) or paclitaxel (B) were serially diluted two-fold (concentrations indicated in the figure). The drug interactions were calculated using the Loewe model on the Combenefit software. Cell viability (% compared to DMSO-treated cells, left panel), synergy matrix as calculated using the Loewe excess model (middle panel), and isobolograms (right panel) for each drug pair are indicated. Presented data are representative of three independent experiments.

**Supplementary Methods.**

**Human samples and immunohistochemistry.** Experiments were conducted in accordance with Bioethics Law No. 2004–800 and the Ethics Charter from the French National Institute of Cancer (INCa), and after approval from the ethics committee of our Institution. PRMT1 immunohistochemistry (IHC) was carried out on tissue microarrays (TMA) containing alcohol, formalin and acetic acid (AFA)-fixed paraffin-embedded tissue, as previously described (2,3). Endogenous activity was blocked with Dako REAL Peroxidase-Blocking solution (DAKO) followed by incubation with rabbit PRMT1 polyclonal antibody (Table S1). This antibody recognizes residues 298-318 within the C-terminal domain of all isoforms of PRMT1. For each tumor sample, proportion of positive cells and its corresponding immunostaining intensity was defined, allowing determination of a score that was defined by the following formula: % of PRMT1-positive tumor cells (%) x intensity of immunostaining. For surface staining quantifications, whole digital slide images were obtained using virtual microscopy (Philips Ultra Fast Scanner 1.6 RA) and analyzed with Digital Image Analysis platform HALO (version 3.0.311.218; Indica Labs). Tissue classifier was trained to segment tumor tissue and stroma. Area Quantification module (v2.1.3) was used to evaluate the area of each tissue class and the area of tissue positive for PRMT1 staining. PRMT1 antibody was validated and optimized for IHC using AFA-fixed cell pellets from MDA-MB-468 treated with PRMT1 siRNAs or control siRNA for 72h (Sup. Fig 1D).

**Colony formation assays.** Cells transfected with siRNA were seeded in six-well plates in 2 mL of growth media. Cells were incubated at 37°C for 6 mitotic cycles (6-14 days), depending on the cell line, until colony formation. Colonies were fixed and stained with 500µL of coomassie blue solution for 20 min. Colonies were photographed by using a LAS-3000 Luminescent Image analyser (Fuji, FSVT) or Chemidoc MP imager (Biorad) and quantified by ImageJ 1.43u software (NIH).

**Soft agar assay.** For soft-agar colony formation, 0.35% agar containing siRNA transfected MDA-MB-468 cells was overlaid onto precast 0.5% bottom agar and incubated 4 weeks as described (4).

**Chromatin Immunoprecipitation (ChIP).** Chromatin was prepared from $4{\times}10^6$ untreated MDA-MB-468 cells using the simple ChIP plus enzymatic chromatin IP Kit (Cell signaling, #9004), following manufacturer's protocol. The chromatin was immunoprecipitated using an anti-PRMT1 or anti-IgG antibodies (Table S1) overnight and the chromatin/antibody complex was pulled down using protein G agarose beads (provided with the kit). Following different washing steps, the chromatin was eluted, and the cross links were reversed using proteinase K. DNA was purified using the spin columns included in the kit and a qPCR was performed using specific primers designed based on published ChIP-seq dataset for PRMT1 (5) for the promoter region of each gene (Table S1).

**Real-time – quantitative PCR assay (RT-qPCR).** For Wnt target gene expression, MDA-MB-468 cells transfected with siRNA were serum-starved overnight and stimulated with Wnt3a conditioned media at 100ng/mL for 6h. RNA was extracted using the RNeasy Mini Kit (Qiagen, 74106) following the manufacturer's protocol. Reverse-transcription and RT-qPCR were performed in a one-step reaction using the QuantiTect SYBR Green RT-PCR Kit (Qiagen, 204245), according to the manufacturer's protocol. The acquisition was done using QuantStudio™ 12K Flex Real-Time PCR System (Applied Biosystems).

**Wnt/β-catenin-activated reporter (BAR) luciferase assay.** 24h post siRNA transfection, MDA-MB-468 cells were transfected with the SuperTOPflash (7X Wnt response element containing plasmid) and pRL-TK-Renilla plasmids (obtained from Institut de Recherches Servier, France) at a 10:1 ratio using X-tremeGENE™ HP (Sigma-Aldrich, 6366236001) as a transfectant. The cells were serum-starved overnight, i.e., 4-5h post DNA transfection and stimulated with 100ng/mL of Wnt3a conditioned media for 6h. Dual-luciferase assay (Promega, E1910) was performed following manufacturer's protocol and the luminescence signal was measured on the Infinite M200 spectrophotometer (Tecan). The ratio of the signal from firefly (SuperTOPflash) to renilla (pRL-TK-Renilla) luciferase was calculated to obtain normalized luciferase activity, representing Wnt/β-catenin activity.

**Drug combinations.** MDA-MB-468 cells were seeded 48h prior to treatment in a 96-well white transparent bottom plate (Greiner Bio-One, 655098) and treated with varying concentrations of the drugs/inhibitors as indicated in Table S1. Cell viability was determined after 7 days of treatment by CellTiterGlo assay (Promega, G7572). Luminescence signal was measured in a Spark spectrophotometer (Tecan). Drug pair interactions using the Loewe model were calculated on the Combenefit software (6). All drug combinations were performed in triplicate reactions per experiment.

**Mice, treatment, and tumor growth measurements.** Six-week-old female Swiss-nude mice were purchased from Charles River and maintained in specific pathogen-free conditions. Their care and housing were as per institutional guidelines as put forth by the French Ethical Committee. GSK3368715 (ChemScene LLC, CS-0100240) was formulated in 10% DMSO (Sigma-Aldrich) at 80 mg/ml and subsequently diluted in water. GSK3368715 toxicity studies were performed by administration of 100 mg/kg daily, to nude mice.

MDA-MB-468 cells ($12x10^6$ per mouse) were injected subcutaneously into nude mice until tumors reached 70 $mm^3$. The tumor fragments obtained from 2 mice were then grafted into the inter-scapular fat pad of nude mice. Xenografts were randomly assigned to control or treatment groups (n = 6/group) when tumors reached a volume comprised between 60 and 80 $mm^3$ and treated with vehicle or GSK3368715 at 80mg/kg once daily orally 5 days/week. During the weekends, the inhibitor was added to the drinking water of mice. Tumor volume was evaluated by measuring two perpendicular tumor diameters with a caliper, twice a week. Mice were euthanized after 8 weeks of treatments. Tumor volumes were calculated as V = a × b2/2, a being the largest diameter, b the smallest. Tumor volumes were then reported to the initial volume as Relative Tumor Volume (RTV). Means of RTV in the same treatment group were calculated, and growth curves were established as a function of time.

**References.**
